# Accounting for Defective Viral Genomes in viral consensus genome reconstruction, application to influenza virus

**DOI:** 10.64898/2026.03.11.711002

**Authors:** Kévin Da Silva, Nadia Naffakh, Marie-Anne Rameix-Welti, Frédéric Lemoine

## Abstract

In the context of viral epidemic surveillance, generating accurate consensus viral genomes from sequencing data is critical for tracking the emergence of mutations of concern, evaluating the genomic diversity of circulating viruses, and anticipating which viral strains could become most prevalent. However, this task is made difficult by the presence of Deletion-containing Viral Genomes (DelVGs), which contain truncated (or rearranged) and potentially mutated versions of the full length virus genome. Because these DelVGs can outnumber the full genome in terms of coverage, potential DelVG specific mutations may be erroneously incorporated into the final consensus, thereby compromising its accuracy. Automatic detection of these DelVGs and of the genomic positions that may harbor DelVG specific mutations is therefore crucial. Here, we present DIPScan, a new method able to (i) accurately and efficiently detect DelVGs in short read datasets, and (ii) mask or correct positions in the consensus genome that may be affected by DelVG-specific mutations. Using several hundreds of simulated and real patient-derived NGS datasets from the National Reference Center (NRC) for respiratory viruses at Institut Pasteur, we demonstrate the capacity of DIPScan to accurately and efficiently detect DelVGs and to correctly adjust the consensus sequences. DIPScan is implemented as a Nextflow workflow, making it highly flexible, scalable, and reproducible, and is now used routinely at the NRC.

## Introduction

Routine virological surveillance has been shown to be of utmost importance for monitoring viral epidemics, especially for (i) monitoring the emergence of mutations of concern (e.g. drug resistance mutations, or other advantageous mutations), (ii) monitoring the genomic diversity of circulating viruses and identifying the main circulating lineages, and (iii) predicting the phylogenetic clades that may be most prevalent in the near future.

Regarding these three topics, high throughput sequencing associated with efficient bioinformatics methods have been the cornerstone of virological surveillance in the past 15 years, and many efforts have been made to improve these methods.

Since the COVID-19 pandemics, from the bioinformatics point of view, many data analysis workflows have been developed and used routinely in surveillance laboratories to process virus sequencing data on a large scale and produce consensus sequences for the main circulating respiratory viruses (e.g. SARS-CoV-2, Influenza, RSV) and other non respiratory viruses (e.g. Ebola, Zika, etc.). Among these workflows, each with its unique aspects depending on the type of virus and data, notable examples include Viralrecon [1, 2], wf-flu (https://github.com/epi2me-labs/wf-flu), ViReflow [3], ViralFlow [4], SARS2seq (https://github.com/RIVM-bioinformatics/SARS2seq), IRMA [5], ARTIC fieldbioinformatics (https://github.com/artic-network/fieldbioinformatics), to name a few.

These workflows usually work as follows: (i) They map the sequencing reads against a reference genome (the closest to the input samples) using a read mapper (e.g. bwa [6]), (ii) They look for variations between the sequencing reads and the reference genome (SNP calling) using tools such as Samtools [7] and iVAR [8], (iii) They modify the reference according to the observed differences to obtain a consensus genome (e.g. using iVAR), in which each position consists of the nucleotide present in roughly more than half of the mapped reads at this position. Variants of this simple workflow exist, with additional steps such as reference selection, assembly, filtering, etc. Each step of these workflows is critical to build a reliable consensus sequence [9]. For example, selecting the right reference is of paramount importance, especially for very diverse viruses such as influenza viruses. Choosing the right parameters for SNP calling and consensus building also has a strong influence on the final consensus, such as minimal acceptable coverage, ambiguous nucleotide assignment, regions of the reference genome to mask, etc.

However, one often-overlooked characteristics of the samples can affect the reconstruction of a high-quality consensus sequence is the presence of Deletion-containing Viral Genomes (DelVGs) in the analyzed samples, that can produce defective interfering particles (DIPs) [10].

### Definition and types of DIPs

DIPs have been initially discovered several decades ago in Influenza virus as non-replicative incomplete viruses [11] that can interfere with the replication of the non-defective homologous virus [12]. Initially, DIPs were thought to be primarily generated in cultured cells and considered insignificant in natural infections [13]. DIPs have since been detected in various virus families, including DNA viruses, RNA viruses, and retroviruses [14–16].

They consist of “viral particles that contain normal structural proteins but only a part of the viral genome” [14]. DIPs are generated during replication and are incapable of replicating independently, requiring the presence of a full-length (helper) virus to complement the missing sequences and corresponding proteins [15].

According to the definition of Vignuzzi and López [14], defective viral genomes (DVGs) can be classified in three main categories depending on how they were generated: (i) DVGs resulting from mutations in the viral genome (single mutation, frameshifts, or hypermutations); (ii) DVGs resulting from large deletions or rearrangements (i.e. DelVGs, the focus of this study), which are thought to form when the viral polymerase detaches from the template strand and resumes replication at a distant point, skipping an large internal genomic region; and (iii) DVGs resulting from snap-back or copy-back, resulting in the duplication of a sequence in reverse complement.

### Mechanisms of production and role of DelVGs

In influenza virus DelVGs, deletions are mainly found in the polymerase segments (PB1, PB2, PA). This was previously thought to be related to the greater length of these segments, but this has not been conclusively confirmed [10]. Moreover, deletion hotspots have been shown to be located at the segments’ termini, within the first 400 nucleotides after initiation, for reasons that are not totally elucidated so far, but maybe related to the presence of A/U rich sequences, direct repeats, or to the formation of a t-loop RNA structure, although with no specific sequence motif [10, 17–19].

While it has been hypothesised that internal deletions can occur through homologous recombination in positive-strand RNA viruses, such recombination events are rare in negative-strand viruses such as influenza viruses. Recent studies suggest that DelVGs production may be a process regulated by viral factors or host factors.

Regarding viral factors, the structure of viral ribonucleoproteins, which could influence break and rejoin sites [15, 20], and, possibly, mutations on the influenza polymerase [21], may be linked to deletions. Also, specific sequences in the RSV genome have been shown to regulate the formation of certain DVG types [22]. Regarding host factors, some cell types seem to produce low amount of DIPs [23, 24].

DVGs have been proposed to have several key roles. Their primary characteristic is the ability, upon co-infection with a full length virus, to interfere with its replication by competitive inhibition and to accumulate while the proportion of non-defective particles decreases, leading to the term “Defective Interfering Particles” [25]. Beyond this, they have been suggested to promote viral persistence in vitro, to modulate the host innate immune response [26, 27], and to contribute to viral evolution by providing a diverse sequence pool for recombination or reassortment with the standard genome [14, 16].

The significance of defective viral genomes is also well-established in influenza virus research more specifically. First, there is accumulating evidence that these defective genomes influence disease severity, and may contribute to viral persistence [24, 28].

Vasilijevic *et al.* correlated a low abundance of DIPs in patients infected with influenza virus with severe disease outcomes [21], while Penn *et al.* suggested that DIP levels during infection influence H5N1 pathogenesis in mice [29].

These properties have prompted the consideration of defective viral genomes as antivirals [30–32] or vaccine adjuvants [14].

### How DelVGs impair consensus reconstruction

A standard method for reconstructing consensus viral genomes from sequencing data involves mapping sequencing reads to a reference sequence and adjusting the reference sequence based on observed variations. However, this approach can be problematic when defective viral genomes, especially DelVGs, predominate in the sample over full-length genomes. In such cases, read mapping (as shown in Fig. 1.B) reveals high coverage at the reference genome’s termini (present in both defective and full-length genomes) but low coverage in the central region (present only in the minority full-length genome).

**Fig 1.**
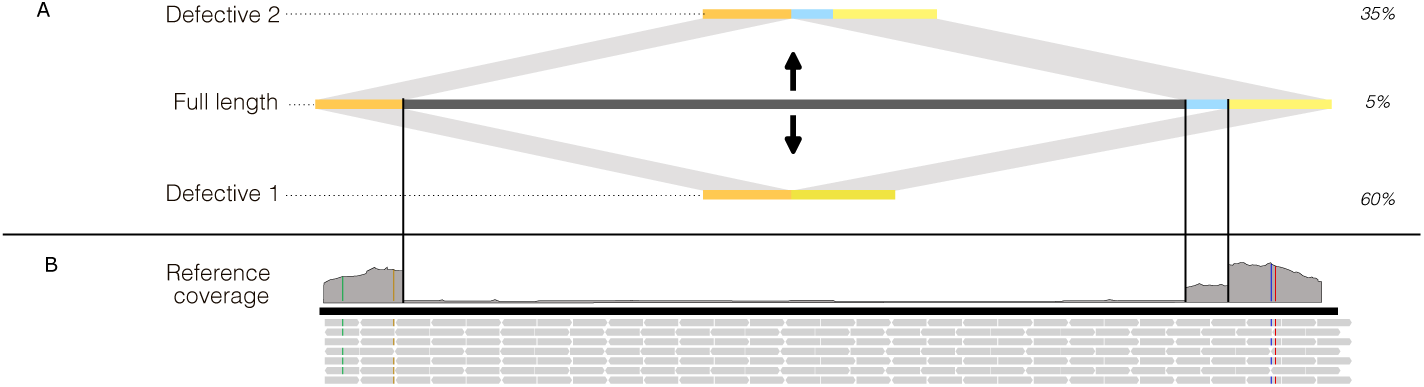
Challenge posed by defective sequences. This schematic illustrates a virtual sample containing two distinct defective sequences, in addition to the full length sequence. **A.** The composition of the two DelVGs, each resulting from the deletion of a unique genomic region. In this virtual scenario, *DelVG 1* constitutes the majority at 60% of the sample sequences, *DelVG 2* accounts for 35%, and the full-length, functional sequence, represents the remaining 5%. **B.** Alignment of sequencing reads to the reference genome, and the resulting coverage profile. Elevated coverage at the termini, accompanied by sharp drops (one near the 3’ end and two near the 5’ end), indicates the presence of two types of DelVGs. Due to the substantial overrepresentation of DelVGs compared to the full-length genome, mutation frequencies at the extremities must be interpreted with caution, as they likely reflect variants in the defective sequences rather than in the full-length genome.

Furthermore, defective particles may carry unique mutations that are absent in the full length genome, as well as in the reference genome. Since these mutations appear in most reads, they are often incorrectly incorporated into the final consensus genome.

Overlooking DelVGs during consensus reconstruction can thus lead to a chimeric sequence that misrepresents the true viral population in the sample. The improper integration of mutations may introduce severe errors, such as premature stop codons, or frameshifts. Fig. 1 illustrates this challenge, depicting how DelVGs with distinctive mutations can distort the reconstructed consensus genome when present in sequencing data.

One possible approach to addressing DelVGs in sequencing is to manually review read alignments and exclude affected segments (in segmented viruses like influenza) or discard entire samples (in non-segmented viruses). However, this method is labor-intensive and poses the risk of losing important surveillance data. Some opt to retain the original consensus sequence under the assumption that DelVGs are rare and negligible in clinical samples. Yet, as demonstrated by Saira *et al.* [33], defective genomes frequently appear in Influenza A/H1N1pdm patient samples (13 over the 26 studied), making their proper handling essential.

### Existing tools to study DIPs/DelVGs

The generation of accurate consensus sequences necessitates the automated detection of DelVGs in sequencing data. This comes down as identifying and correcting problematic sites (i) where read coverage is higher in the defective genome relative to the full-length genome, and (ii) that have mutations specific to the defective genome.

Several tools, such as DI-TECTOR [34], ViReMa [35, 36], DG-Seq [19], DVG-profiler [37], VODKA [22], or VODKA2 [38] have been developed over the past decade to identify defective genomes in sequencing datasets. However, while these tools can detect various types of defective genomes, (i) they are not suited for large-scale, automated consensus construction in routine settings, and (ii) recent studies have demonstrated low agreement between them when identifying deletion junctions [39].

DI-TECTOR relies on bioinformatics tools that are not adapted to current data production scales, making it impractical for modern dataset sizes. Moreover it is designed primarily for defective genome detection rather than the generation of a clean final consensus genome. ViReMa was developed to detect recombination events in general. While it is designed to map split reads and therefore detect large deletions, it is not designed to clean or correct the full length consensus sequence. DVG-profiler identifies reads that do not align perfectly to the reference. For each read with multiple alignments, it tries to pair alignments that could represent two segments of a single DelVG read. Orientation of the alignment pair determines the type of defective genome, and the number of reads spanning a junction compared to the average full-length coverage determines the defective/full-length ratio. Using only junction-spanning reads may underestimate the proportion of defective genome. Additionally, it is not designed to clean or correct the full length consensus sequence present in the sample. DG-Seq was developed to detect and quantify influenza defective genomes, but not to correct the full length consensus. As for VODKA, it was developed to specifically identify copy back defective genomes, which are not our primary focus, occur less frequently, and make the detection of interesting regions to correct more difficult. Its update, VODKA2, increases its accuracy and speed, and expands its functionality to the detection of delVGs. However, VODKA and VODKA2 are designed to detect and count junction reads, they do not provide DelVG quantification nor full-length consensus correction.

Here, we introduce DIPScan, a bioinformatics workflow specifically designed to identify deletion-containing viral genomes (DelVGs) resulting from large deletions within Illumina sequencing datasets and, when necessary, correct consensus sequences by introducing the likely nucleotide of the full length genome or by masking potentially problematic positions at these positions.

DIPScan requires the following inputs: (i) an Illumina sequencing read file in FASTQ format, (ii) a reference genome in FASTA format, and (iii) an optional precomputed consensus sequence also in FASTA format. The workflow generates two key outputs: (i) a report indicating the presence or absence of DelVGs along with the proportion of of internal deletions for each genomic segment, and (ii) a corrected consensus sequence.

## Material and Methods

### DIPScan workflow

DIPScan features seven primary computational steps shown in Fig. 2 and described below: mapping, snp calling, extraction of deletion boundaries, metrics computation, breakpoints selection, DelVGs proportion estimation, and consensus correction.

**Fig 2.**
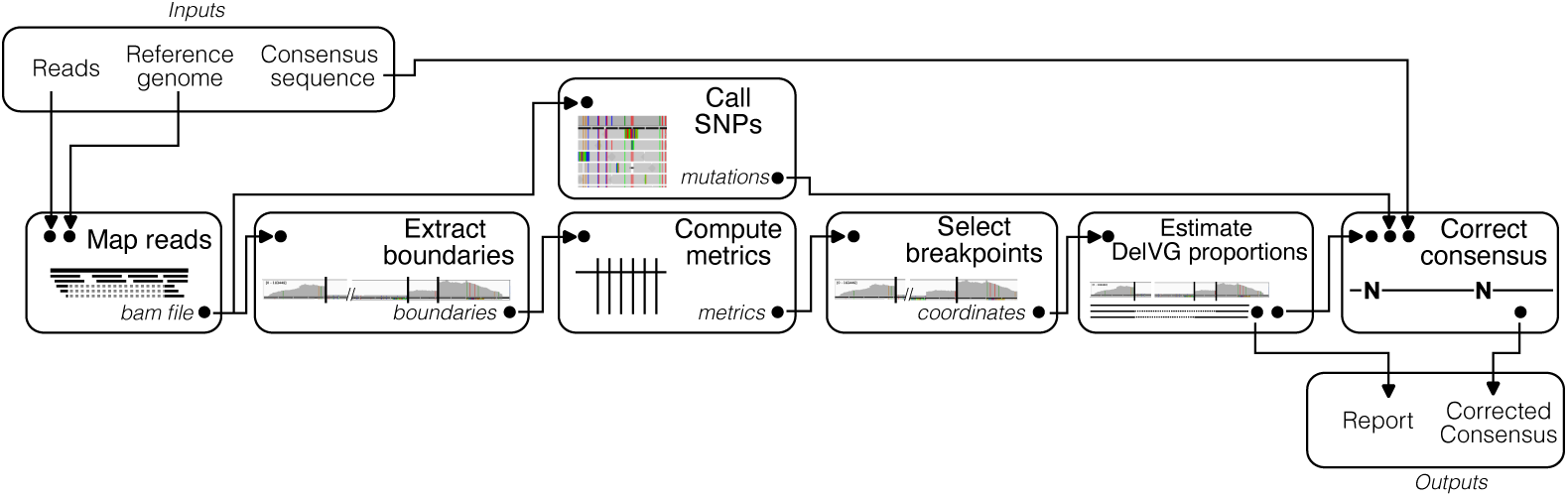
DIPScan workflow. A simplified schematic representation of the DIPScan workflow for identifying DelVGs and correcting the consensus sequence. The pipeline requires an input read file, a reference genome, and an optional user-supplied consensus sequence. Processing consists in seven main steps: (i) read mapping, (ii) SNP calling, (ii) boundary extraction, (iv) metric computation, (v) breakpoint selection, (vi) DelVG proportion estimation, and (vii) consensus sequence correction (if an input consensus sequence is provided). The final output includes two files: a report of identified DelVG boundaries, and the corrected consensus sequence.

#### Mapping split reads

DIPScan begins by mapping the sequencing reads to the user-provided reference genome. This is achieved using a two-step approach that accounts for split reads spanning large deletions. In the first step, the reads are mapped to the reference genome using BWA-MEM2 [40]. From the resulting alignment, only the reads that map with at most 8 clipped nucleotides (unmatched read extremities) are kept. This produces a classical alignment, containing reads that map perfectly, as well as those containing SNPs or short insertions/deletions and less than 8 clipped positions.

In the second step, all unmapped reads and those that map with at least 8 clipped nucleotides are extracted, as they can define split regions in the case of DelVG. These reads then undergo STAR alignment [41] (Spliced Transcripts Alignment to a Reference, v2.7.11a). Although STAR was originally designed for aligning RNA-Seq reads across splicing sites, it is also highly effective at accurately mapping split reads flanking large deletions (with specific run parameters, see S1 Text), with read portions on either side of the deleted sequence, that are considered as “non-canonical splice sites”. This makes STAR well-suited to identifying defective genomes of interest, as these split reads often indicate large-scale structural variations within the genome, such as deletions, insertions, duplications, or translocations.

The outputs of these two steps are merged together in a single mapping file (BAM format) containing full reads mappings and split read mappings.

#### Extraction of deletion boundaries

The mapping file is then used to identify potential deletion boundaries by extracting reads containing “skipped regions” (denoted by ’N’ in the CIGAR string of the SAM format) of more than 150 nucleotides. These “skipped reads” each span deleted regions, defining start and end coordinates relative to the reference genome. The resulting position pairs are consolidated into a non-redundant set for downstream analysis.

#### Computing defective metrics

Using this set of unique start-end position pairs, various metrics are calculated. For each pair, these metrics include:

- *Supporting reads*: The number of reads spanning the exact deletion coordinates.
- *Split frequency* : The proportion of *supporting reads* relative to the total number of split reads (at any deletion coordinate). This value indicates the relative frequency of this particular deletion compared to other deletions.
- *Total frequency* : The proportion of *supporting reads* relative to the total number of mapped reads. This value indicates the relative frequency of this particular deletion compared to the whole sequencing dataset.
- *Expected minimum frequency* : The expected *Total frequency* if the proportion of DelVG was 2%. This value corresponds to 2% of read length divided by the non-deleted region size (reference genome size minus deletion length).
- *Deletion start ratio*: The local deletion-start prevalence at a breakpoint is quantified as the ratio of mean coverage over the 5 nucleotides downstream to the mean coverage over the 5 nucleotides upstream of the breakpoint start.
- *Deletion end ratio*: The local deletion-end prevalence at a breakpoint is quantified as the ratio of mean coverage over the 5 nucleotides upstream to the mean coverage over the 5 nucleotides downstream of the breakpoint end.

#### Breakpoint selection

In order to filter out lowly prevalent breakpoints that may add noise in the data, the following default filters (though adjustable depending on the sequencing depth) are applied to ensure the robustness of detected breakpoints:

- *Supporting reads* should be greater than 100 reads. This criterion eliminates background noise.
- *Total frequency* should be greater than the *Expected minimum frequency*. This removes breakpoints that are poorly represented relative to the entire dataset.
- Both *Deletion start ratio* and *Deletion end ratio* must be smaller than 1. This removes meaningless breakpoints having inner-coverage higher than outer-coverage.

The filtered list may include multiple breakpoints that delineate various deleted regions and, notably, several DelVG sequences.

#### DelVG proportion estimation

The potential risk posed by DelVGs in viral consensus reconstruction is particularly significant when DelVGs are dominant relative to the full-length genome. In such cases, a potential mutation may become dominant and subsequently incorporated into the final consensus sequence. Estimating the proportion of each potential DelVG is therefore important.

For a given segment, the *n* previously selected unique breakpoint positions (either starts or ends) divide the segment into *n* + 1 distinct regions. The median nucleotide coverages, denoted as *cov*_1_*, . . . , cov_n_*_+1_, are computed for all the regions *r*_1_*, . . . , r_n_*_+1_. Because coverage frequently declines at the edges of the first and last regions, the median coverage is calculated after excluding the first (or last) 150 nucleotides of these regions, respectively. If the first (or last) region is shorter than 150 nucleotides, the median coverage of the last (or first) region is used instead. If both regions are shorter than 150 nucleotides, the higher coverage value between the two is selected.

Given that the *n* breakpoint positions represent *d* distinct DelVGs (some breakpoints starts or ends may be shared across multiple DelVGs), our objective is to estimate the relative abundances *{c*_1_*, …, c_d_, c_d_*_+1_*}* of each DelVG (1*, . . . , d*) and of the full-length segment (*d* + 1) in the sample. These estimates should best explain the observed coverages *cov*_1_*, . . . , cov_n_*_+1_ across the *n* + 1 regions.

This estimation is formulated as a system of *n* + 1 linear equations, where each equation models the coverage *c_r_* of a region *r* as a weighted sum of the contributions from all sequences (DelVGs and the full-length segment):

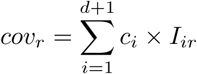

Here, *I_ir_*= 1 if region *r* is present in sequence *i*, and 0 if deleted, while for the full length sequence (*i* = *d* + 1), *I_ir_*= 1 for all regions.

Beyond coverage of all regions, we also leverage split-read counts at each breakpoint, where each split-read count *s_i_* represents the number of reads spanning the breakpoint present in DelVG *i*. Using these split-reads counts, we calculate the relative proportion *p_i_* of each DelVG compared to all DelVGs:

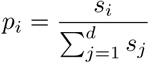

To ensure consistency, the estimated DelVG abundances *c*_1_*, . . . , c_d_* should also satisfy:

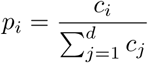

This constraint can easily be reformulated as:

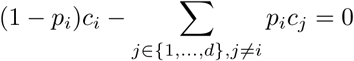

yelding *d* additional equations that are integrated into the system. An illustrative example with 3 DelVGs is given in S1 Fig. Because these equations have no exact solution (no abundance coefficients satisfy all constraints), we estimate the sequence abundances that best match the data using Non-Negative Least Squares (NNLS) [42], as implemented in Scipy (v1.15.3).

After solving, the abundance coefficients are normalized by their sum to obtain the proportions of each DelVG and the full-length segment.

A sample is flagged as containing relevant DelVGs if the sum of DelVG proportions (excluding the full length segment) exceeds 50%.

#### Consensus correction

Consensus correction relies on a predefined list of candidate mutations. To generate this list, DIPScan first identifies potential variant positions relative to the reference genome using iVar (v1.3) [8], applied to a filtered BAM file from which noisy split reads (those not mapping to selected breakpoints) have been removed. This produces a tabulated output containing all candidate mutation sites, along with comprehensive details for each, such as the mutation type (SNP, INDEL, reference/alternative nucleotides), the number of reads supporting the alternative allele, the total read coverage at the position, the statistical confidence metrics for the mutation, etc. This information then feeds into the consensus correction pipeline, which comprises the following steps, as illustrated in Fig. 3:

**1) Identification of regions to be corrected** The breakpoints identified previously are used to define the maximal region that may harbor dominant DelVG mutations and therefore necessitates correction. This maximal region is composed of two parts:

1. The *start region* (orange sequence on Fig. 3): spanning from position 0 to the right-most position among the start positions of the breakpoints.
2. The *end region* (blue sequence on Fig. 3): extending from the left-most position among the end positions of the breakpoints to the end of the reference segment.
**2) Selection of mutated positions** DIPScan selects positions located within either the start or the end region of the genome (in red, marked with a star in Fig. 3) that may need correction. To do so, the initial consensus sequence is aligned against the reference genome using MAFFT [43] (v7.525 -auto) in order to match the positions of the user-provided consensus with the positions of the reference genome. Then, using the alignment along with the called mutations from iVAR output, DIPScan selects positions that are covered by at least 10 reads, and specifically for the positions of the consensus with SNPs, also those that are affected by a statistically significant mutation (with a *p.value ≤* 0.05), either major or minor, as computed by iVAR.
**3) Choosing the right nucleotide** At this final step, for a given mutated position, we aim at associating each observed nucleotide to one or several sequences (DelVGs and full length), and use this information to incorporate the nucleotide associated to the full length sequence in the consensus. This can be done by leveraging the estimated frequency of each sequence (DelVG and full length), and the frequency of each nucleotide at the given position. But first, we can already resolve straightforward cases:

1. If the estimated proportion of the full-length genome is below a mutation threshold *m_t_* (*p_fl_ ≤* 1 *− mt*, by default *m_t_*= 0.98), then the genome is considered too low in frequency to confidently determine the correct nucleotide at any position called by iVAR, and an N is incorporated at all the mutated positions in the consensus.
2. If the mutation frequency *f_mi_* falls within the ambiguous range *m_ar_* (by default, *m_ar_* = [0.45, 0.55]), the mutation cannot be reliably identified, and an N is incorporated in the consensus.
3. If 0.55 *< f_mi_ < m_t_* (major mutation) and *p_fl_ ≥* 0.5 (the full length sequence is dominant), then the mutation is retained in the consensus.
4. If 1 *− m_t_ < f_mi_ <* 0.45 (minor mutation) and *p_fl_ <* 0.5 (the full length sequence in not dominant) and the frequencies of full-length (*p_fl_*) and DelVG (*p_dip_*) variants do not overlap (*i.e. |p_fl_− p_dip_| > δ*, with *δ* = 0.1 by default) and the frequency of the mutation, *f_mi_*, is compatible with the full length frequency *p_fl_* (*i.e. |p_fl_− f_mi_| <*= *δ*, with *δ* = 0.1 by default), then the minor mutation is incorporated in the consensus sequence.

**Fig 3.**
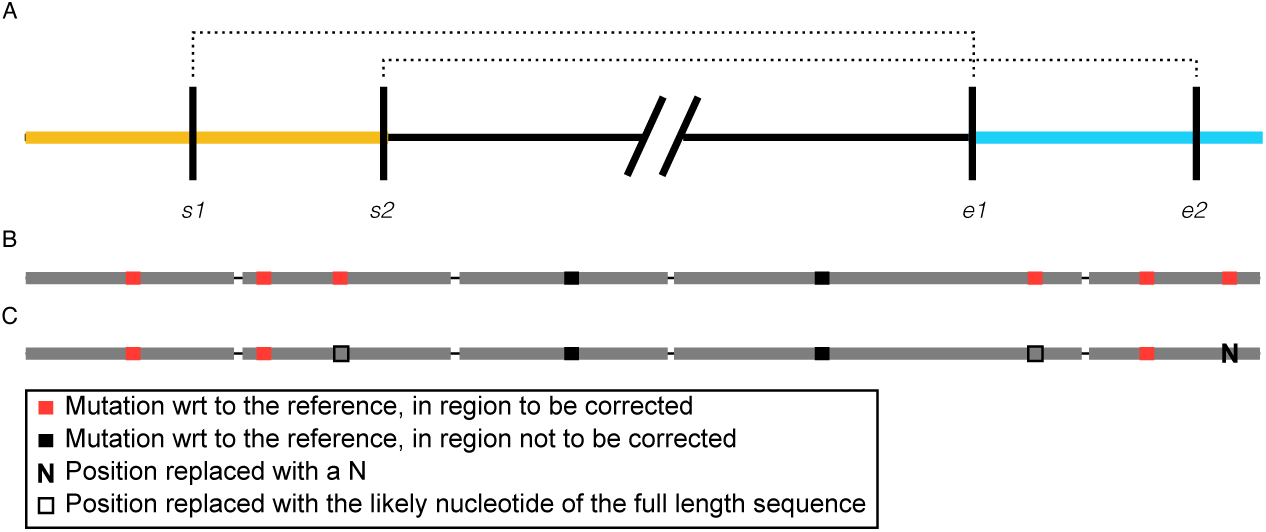
Schematic representation of the consensus sequence correction process. **A.** Defining regions to be corrected. Based on two previously identified breakpoints (coordinates [*s*1*, e*1] and [*s*2*, e*2]), two regions are flagged for potential correction: the start region (in orange) spans from coordinate 0 to the rightmost breakpoint start (*s*2), and the end region (blue) spans from the leftmost breakpoint end (*e*1) to the end of the reference genome. The central sequence (in black), with coordinates [*s*2*, e*1] is protected from correction as it is presumed to come exclusively from reads originating from the full length genome. **B.** Mutation detection. The initial consensus sequence is aligned against the reference genome. Mutations that fall within the designated correction regions are identified (highlighted in red). **C.** Correction of the identified positions in the consensus. Each identified mutation is resolved through one of the following ways: (i) replacement with an ambiguous character ’N’, (ii) retention of the mutation when it is presumed to originate from the full length, or (iii) substitution with the probable full-length nucleotide when the mutation is attributed to a DelVG, the latter determined by comparing its frequency to the coverage of the DelVG sequence.

All other cases correspond to more complex scenarios, such as the presence of multiple DelVGs along with the full length sequence, where sequence frequencies must be combined to match to the frequency of a given nucleotide. In this scenario, for a given mutated position, we try to match the detected sequences (DelVGs and full-length) with the possible nucleotides found in the reads at this position, such as the sum of the sequence frequencies associated to a given nucleotide is approximately equal to the frequency of the nucleotide. This corresponds to a variant of a well-characterized problem, known as the “Multiple Subset Sum”, and can be formulated as “For each label (nucleotide), identify a subset of objects (sequences) whose combined frequencies match the label’s frequency”. While this is an NP-hard problem, heuristics (such as simulated annealing or MCMC) can be used to find solutions. Moreover, since the number of sequences and unique nucleotides is very small in our case, a solution is findable very quickly. The solution is then used to assess whether the full length sequence can be associated unambiguously with one nucleotide (the nucleotide is then incorporated in the consensus) or not (a N is incorporated in the consensus).

Additionally, to prevent misaligned reads from being mistaken for mutations, we compare the read depth at each potential mutation site with the expected depth accounting for the full-length genome and all defective genomes spanning the position). If the mutation is in a lowly covered region compared to the expectation (the ratio of the position depth and the expected depth is less than 80%), then the mutation is considered unreliable and an N is incorporated.

### Testing datasets

DIPScan was tested on two datasets: (i) A dataset composed of 110 simulated influenza virus sequencing data corresponding to several scenarios, and (ii) a dataset composed of 551 routine sequencing of influenza samples from the NRC for Respiratory Viruses at Institut Pasteur.

#### Simulated dataset

To evaluate DIPScan in a controlled setting, we simulated sequencing datasets containing DelVGs (see Fig. 4). We started with 11 influenza consensus genomes (including all segments) from the real influenza dataset described in the next section, representing various subtypes. These genomes are referred to as *Full Length True Genomes*. We then randomly removed internal regions from one segment (either PB1, PB2, or PA) of these genomes, to create *DelVG True genomes*. Next, we introduced between 1 and 10 new mutations into each *DelVG True genome*, making them distinct from their associated *Full Length True Genome*. These modified genomes are called *DelVG mutated Genomes*. For each *Full Length True Genome* and its corresponding *DelVG Mutated Genome*, we simulated 10 datasets using ReSeq [44] (command reseq illuminaPE –maxFragLen 500) with a full-length/DelVG ratio (based on average coverage per position) ranging from 100% to 10%. This approach allowed us to test both the sensitivity and the precision of DIPScan. This resulted in 110 datasets, further described on S2 Text and S1 Table.

**Fig 4.**
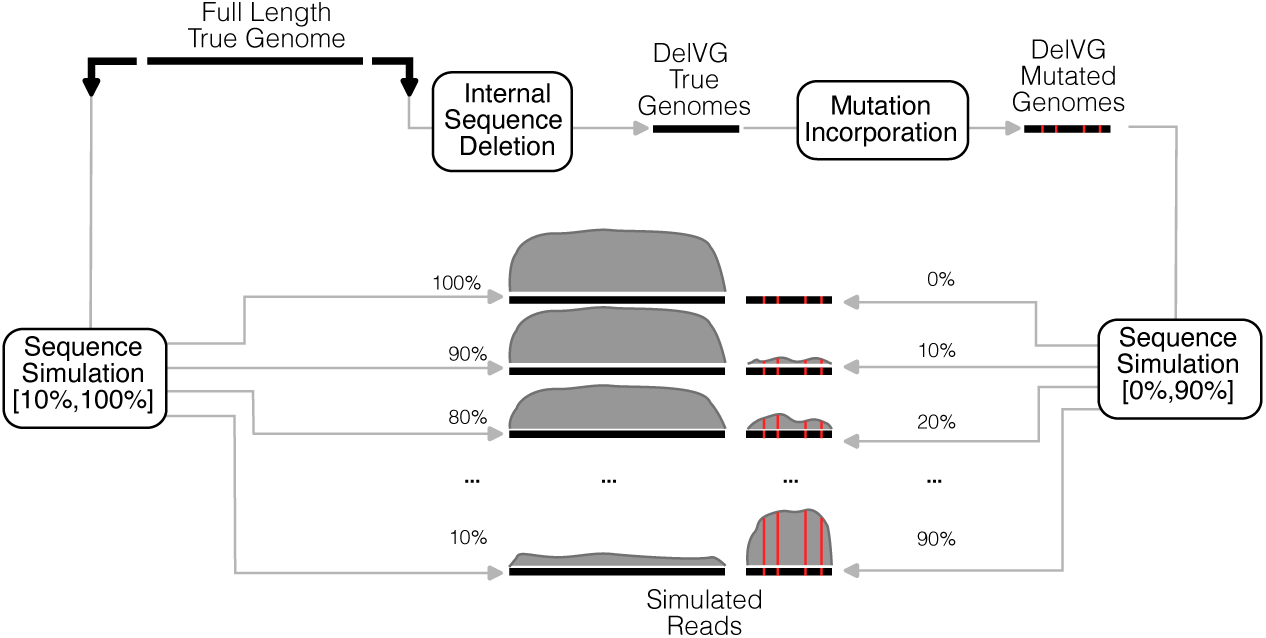
Simulation workflow. Schematic illustration of the simulation workflow for generating mixed full-length and defective genome samples. Starting from a known full-length genome (with 8 segments), an internal sequence of one segment is deleted to create a DelVG sequence, and 1 to 10 random mutations are introduced. Sequencing reads are then simulated from mixtures of the full-length and DelVG genomes at nine distinct ratios(from 100%:0% to 10%-90% full-length:DelVG), yielding ten simulated samples per starting consensus genome.

#### Real influenza virus dataset

We applied DIPScan on 551 real Influenza virus sequencing dataset sequenced at Institut Pasteur’s NRC. It comprises 551 samples collected between 24/02/2025 and 04/04/2025, and consisting of: 125 A/H1N1pdm (22.7%), 232 A/H3N2 (42.1%), and 194 B/Victoria (35.2%). All these samples were visually inspected, to search for DelVG patterns (high coverage on the extremities and low coverage in the central region).

Among these, 55 A/H1N1pdm (44%), 74 A/H3N2 (31.9%) and 103 B/Victoria (53.1%) showed patterns typical of DelVGs resulting from large deletion.

## Results

### **A.** Overview of the prevalence of DelVG sequences in samples

In large-scale sequencing efforts, which often process thousands of samples annually, DelVGs can introduce errors into consensus sequences, leading to their erroneous inclusion in public databases. For instance, during the 2023–2024 season, the National Reference Center (NRC) for Respiratory Viruses at Institut Pasteur Paris sequenced 1,767 influenza samples, of which 540 (30.6%) contained one or more defective segments upon manual review. When restricting the analysis to high-quality sequences eligible for database submission, 471 out of 1,529 samples (30.8%) still exhibited defective segments. A subtype-specific breakdown revealed that defective segments were present in 31.8% of A/H1N1pdm, 30.6% of A/H3N2, and 21.2% of B/Victoria samples, proportions that remained consistent (32.2%, 30.2%, and 21.8%, respectively) when considering only high-quality sequences. Further examination at the segment level demonstrated variability in DelVG prevalence: PB2 had the highest rate (20.1% of high-quality samples), followed by PA (17.7%) and PB1 (14.6%), while HA and NP showed very low rates (0.07% each). No DelVGs were observed in the MP, NA, or NS segments.

These findings underscore the critical need for automated DelVGs detection tools, such as DIPScan, to ensure the accuracy of produced consensus sequences.

### **B.** Simulated datasets

To evaluate the capacity of DIPScan to accurately detect DelVGs and correct the reconstructed consensus genome, we first applied it on the simulated dataset described in the methods section. This dataset consists in 110 virtual influenza virus sequencing datasets, each comprising a mix of reads simulated from one of the 11 original consensus genomes and reads from a corresponding mutated, partially deleted genome segment, with proportions varying from 0 to 90%.

We executed DIPScan on each of the 110 datasets using the default options and assessed its capacity to detect DelVG breakpoints along their proportion, and to correct the consensus sequence. To do so, we computed the following metrics: (i) the detected breakpoints compared to the expected ones, (ii) the estimated DelVG proportion compared to the known proportion in each sample, and (iii) the number of mutations effectively masked or corrected in the final consensus.

#### Breakpoint detection and tool comparison

The first aspect of DIPScan we assessed was its ability to precisely identify DelVG breakpoints within input datasets. Since ViReMa, DG-Seq, and VODKA2 are specifically designed to detect these breakpoints, rather than estimate DelVG proportions or refine consensus sequences, this presents an ideal opportunity to benchmark DIPScan’s performance against these tools. For this comparison, we applied DIPScan, ViReMa, VODKA2, and DG-Seq to each simulated dataset and assessed their performance by matching the identified breakpoints with the ground-truth breakpoints, i.e. those predefined in the simulations. We ran all tools with their default parameters and analyzed their outputs without further processing. Consequently, our analyses incorporated each tool’s built-in filtering criteria. For example, DIPScan by default only reports breakpoints separated by more than 150 nucleotides. Hereafter, breakpoints reported by the tools that coincide with simulated breakpoints are considered true positives; reported breakpoints without simulated matches are considered false positives; and simulated breakpoints absent from tool outputs are considered false negatives.

As shown in Fig. 5, ViReMa, DG-Seq, and VODKA2 detected 1,060, 582, and 270 breakpoints, respectively, of which only 46 (4.3%), 99 (17%), and 64 (23.7%) were true positives. In contrast, DIPScan identified 93 breakpoints, all of which were correct, while missing 6 true breakpoints. Among these missed breakpoints, four originated from low-frequency DelVGs (10%), one from a 20% DelVG, and one from a 30% DelVG, likely due to inaccurate DelVG proportion estimates or insufficient breakpoint read support, below the detection threshold. These 6 breakpoints do not constitute a problem in terms of consensus correction.

**Fig 5.**
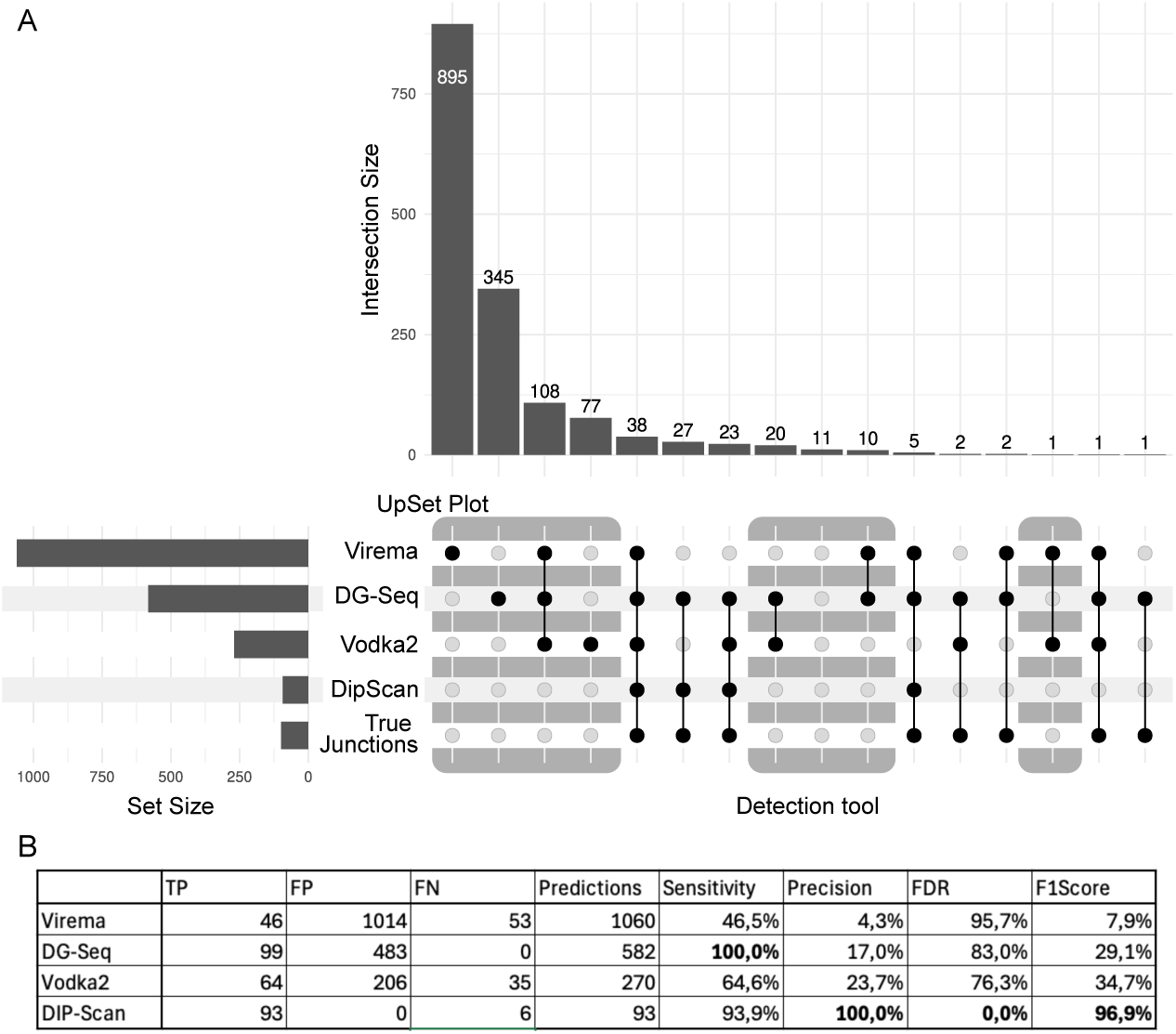
Evaluation of DIPScan for breakpoint detection. Performance evaluation of DIPScan, ViReMa, DG-Seq, and VODKA2 in detecting DelVG breakpoints. (A) UpSet plot illustrating the overlap of breakpoints detected by the three tools and the true ones. For instance, 895 breakpoints are uniquely identified by ViReMa, while 38 breakpoints are correctly detected by all tools. Columns shaded in gray represent false breakpoints. (B) Breakdown of detected breakpoints per tool along the true breakpoints, categorized as true positives (TP), false positives (FP), and false negatives (FN), accompanied by key performance metrics: sensitivity, precision, false discovery rate (FDR), and F1-score.

DG-Seq achieved perfect sensitivity (100%), detecting all true junctions, but at the cost of 483 false positives, resulting in low precision (17%), a high false discovery rate (83%), and an F1-score of 29.1%. By comparison, DIPScan demonstrated superior performance across key metrics, with perfect precision (100%), no false discoveries (FDR: 0%), and a global F1-score of 96.9%.

It is worth noting that all methods except DIPScan incorrectly detected many breakpoints in the 0% DelVG simulation, which contained no true breakpoints. While specific filtering criteria could eliminate many false positive breakpoints, optimization of these thresholds falls outside the scope of this study.

#### DelVG proportion estimation

On the same simulated data, DIPScan accurately estimates the proportion of DelVG sequences. Correlation between the known ratio and the estimated ratio is high (Fig. 6 A., Pearson correlation of 0.99 between average estimated ratios over the 11 samples at each known ratio and the known ratio).

**Fig 6.**
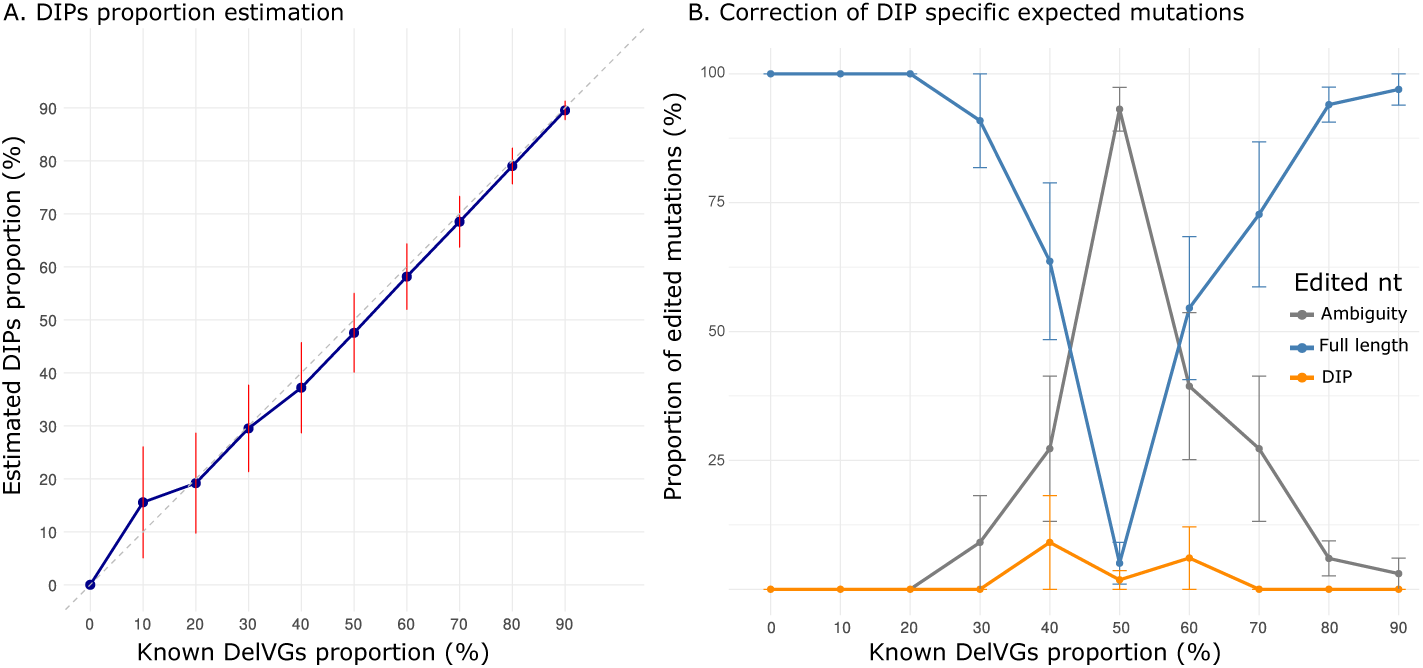
Evaluation of DIPScan for DelVG proportion estimation and consensus correction. **A.** Accuracy of DelVG proportion estimation. The mean distribution and confidence intervals of DelVG ratios estimated by DIPScan (y-axis) is shown for the 11 samples together across each simulated input proportion (x-axis). **B.** Correction performance for DelVG-specific mutations. The proportion of corrected DelVG-specific mutations (y-axis) as a function of the simulated input DelVG portion (x-axis) is represented for the three following outcomes: replacement with an ambiguous base (gray curve), correct reversion to (or retention of) the full-length genome nucleotide (blue curve), or erroneous retention of the DelVG-specific mutation (yellow curve). As expected, ambiguities peak around a 50% DelVG proportion, while correct retention or reversion peaks at both low DelVG proportion (where the full-length genome is dominant) and high DelVG proportion (where the DelVG is dominant, triggering a reversion to the full-length base).

Except for two samples (see S2 Fig), the estimations were slightly under estimated compared to the known proportion, although close to the expectation.

Across all combined datasets, DIPScan demonstrated an overall good accuracy of 94.5% (104/110, six incorrectly classified samples were false negative with low DelVG proportions) in classifying samples as containing DelVGs or not.

#### Mutation detection and consensus correction

In our simulation scenario, three kinds of mutations can in theory be identified relative to the reference genome: (i) Mutations that are shared between the DelVGs and the full length sequence, originating from the original consensus sequence), (ii) mutations that are specific to the DelVGs, randomly introduced, and (iii) mutations that are specific to the full length sequence, if the randomly introduced nucleotide reverts the nucleotide to the reference. The latter case did not happen in our simulations.

As shown in Fig. 6.B and S3 Fig, DIPScan demonstrates a strong performance in handling common and DelVG-specific mutations, respectively.

First, for shared mutations, where no consensus correction is expected, DIPScan demonstrated high fidelity, correctly retaining 99.6% (3976/3980) of the mutations across all samples. Among the 14 changed positions, 11 were masked (replaced by Ns) as these mutations fell just below the default 98% threshold for fixed mutations. The 3 incorrect changes involved samples where the mutations coincidentally displayed the same full-length/DelVG ratio between two nucleotides. This led DIPScan to misclassify them as DelVG-specific mutations.

Then, DIPScan demonstrated good performance in detecting all DelVG-specific mutations introduced into the test samples. Across all samples, it correctly retained or corrected 73.4% (426/580) of full-length mutations, introduced Ns (ambiguous bases) for 23.8% (138/580), and misassigned 2.8% (16/580). Performance varied across the expected DelVG ratio. In the [50%, 90%] range, where DelVG presence complicates consensus reconstruction, DIPScan correctly retained or corrected 59.3% (172/290) of full-length mutations, assigned Ns for 37.9% (110/290), and incorrectly changed 2.8% (8/290). In contrast, for the [0%, 50%[ range, performance improved, with 87.6% (254/290) of mutations correctly retained or corrected, Ns introduced in 9.7% (28/290), and misassignments remaining at 2.8% (8/290).

Notably, all misassigned nucleotides occurred in the same three samples where DelVG proportions were poorly estimated.

Altogether, these results demonstrate the good performances of DIPScan on simulated datasets at detecting DelVGs, their proportion, and correcting the consensus sequences accordingly.

### **C.** Routine influenza virus sequencing datasets

To evaluate DIPScan’s performance in a real-world setting, we applied it to a well-characterized influenza virus dataset that had been manually curated for DelVGs. Manual curation involved visual inspection of genome coverage plots to identify segments with sharp coverage declines, indicating a high probability of DelVG presence.

To do so, we assessed DIPScan based on three criteria: (i) DelVG detection accuracy, (ii) reliability of DelVG proportion estimates, and (iii) capability for consensus sequence correction. Finally, we analyzed the detected DelVG breakpoints to identify potential genomic hotspots.

#### DIPScan detection accuracy

To assess DIPScan detection performances, we classified the results into five categories (Fig. 7):

- **Consistently defective**: both manual detection and DIPScan identified DelVGs in the sample;
- **Consistently not defective**: neither manual detection nor DIPScan identified any DelVGs in the genome;
- **Inconsistent, defective in manual**: DelVGs were identified manually but not by DIPScan. These cases are detailed below, as they may represent either a limitation of DIPScan or manual detection errors.
- **Inconsistent, Defective with DIPScan** (*<* 50%): DIPScan identified DelVGs representing less than 50% of the genomes, which manual detection might miss due to overall coverage similarities between and outside the deletion breakpoints. This category is not considered as errors but rather reflects different strategies in defining DelVGs.
- **Inconsistent, Defective with DIPScan** (*≥* 50%): DIPScan identified DelVGs representing more than 50% of the genomes. This category is detailed below, as it may indicate DIPScan’s oversensitivity or manual detection errors.

**Fig 7.**
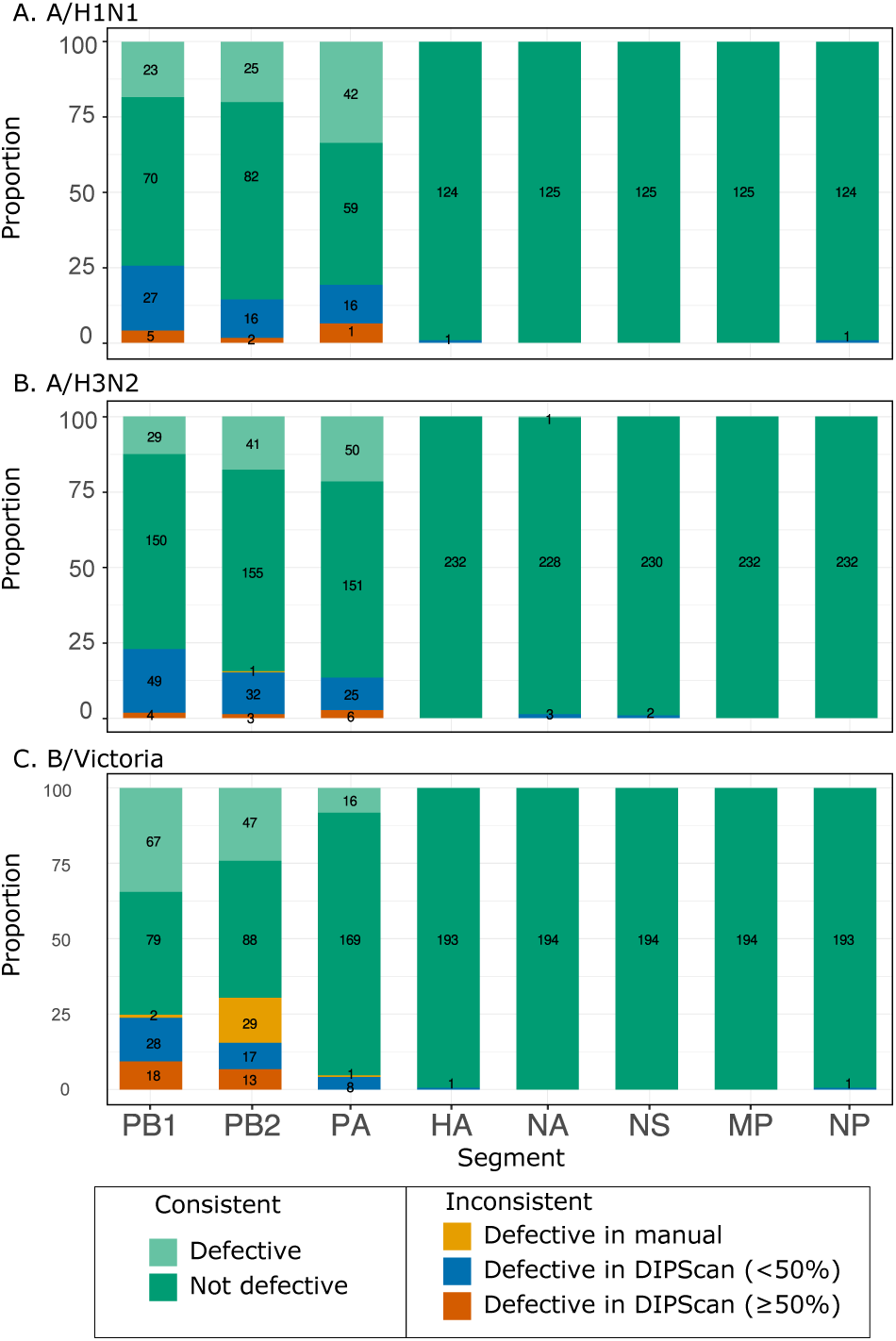
Evaluation of DIPScan on real datasets. Evaluation of DIPScan detection on real Influenza virus sequencing dataset: Manual curation versus DIPScan. Results are categorized based on agreement between methods: Consistent (both methods agree on the presence (light green) or absence (dark green) of a DelVG) or Inconsistent. Inconsistent results are further broken down into: (i) DelVG detected only in manual curation, (ii) DelVG detected only in DIPScan with an estimated proportion below 50%, and (iii) DelVG detected only by DIPScan with a proportion above 50%. The results are grouped by Influenza virus segment and subtype: **A.** A/H1N1pdm (125 samples), **B.** A/H3N2 (232 samples), and **C.** B/Victoria (194 samples). Overall, we observe a good consistency between DIPScan results and manual curation. DelVGs detected only by DIPScan, at a proportion above 50% were usually missed by manual curation.

Across the 4,408 analyzed segments (551 samples), DIPScan and manual detection agreed on 92.8% of segment classifications (Table 1.A). Agreement rates varied by segments: 418 consistent PB1 (75.9%), 438 PB2 (79.5%), 487 PA (88.4%), 549 HA (99.6%), 548 NA (99.5%), 549 NS (99.6%), 551 MP (100%), and 549 NP (99.6%).

**Table 1.** Confusion matrix comparing DIPScan and manual classification of 4,408 segments from 551 samples.

|  |  | DIPScan |  |
| --- | --- | --- | --- |
| A | Manual dataset | Defective | Negative |
|  | Defective | 341 | 33 |
|  | Negative | 286 | 3,748 |
| B | Manual dataset | Defective ($\geq 50\%$) | Negative ($< 50\%$) |
|  | Defective | 341 | 2 |
|  | Negative | 47 | 3,969 |
|  | Unclassifiable | 12 | 37 |

Discrepancies included 33 segments classified as defective only in the manual dataset and 286 detected solely by DIPScan (Table 1.A). Notably, no inconsistencies were found in the HA, NA, NS, NP, or MP segments that could be considered potential errors.

Using manual curation as ground truth, DIPScan achieved 91% sensitivity and 54% precision (92.8% accuracy). However, most disagreements involved DIPScan detecting low-frequency DelVGs (*<* 50%) missed during manual inspection. When considering only high-frequency DelVGs (relevant for consensus reconstruction) as true positives (Table 1.B), DIPScan’s performance improved to 99% sensitivity and 88% precision (98.9% accuracy). Further review showed most remaining discrepancies were either overlooked in manual curation or negligible low-frequency DelVGs, with 49 cases remaining unclassifiable (Table 1.B). These unclassifiable cases presented a complex scenario where it was unclear whether manual review or DIPScan was correct.

In particular, among the cases where DIPScan detected DelVGs with a proportion *≥* 50% but manual review did not (**Inconsistent, Defective with DIPScan (***≥* 50%)), 12 samples belonged to the A/H1N1pdm subtype. Upon re-evaluation, all instances affecting the PB2, PB1, and/or PA segments (*n* = 15) were found to be potential manual errors, primarily due to the difficulty in visually evaluating the proportions of full-length genomes over DelVGs. For the A/H3N2 subtype, 11 samples were identified, with all cases affecting the PB2, PB1, and/or PA segments (*n* = 13) ultimately attributed to potential manual errors. For the B/Victoria subtype, 29 samples were identified, all affecting the PB2, PB1, and PA segments. In 12 cases, the inconsistency was due a complex scenario likely involving a defective genome with a single breakpoint, resulting in truncation of the entire 5’ part. One case was due to a manual report error since a DelVG was unambiguously visible, and 18 cases were found to be also potential manual errors.

#### DelVG proportion estimation

Direct assessment of DIPScan’s accuracy in estimating DelVG proportions in real viral samples is inherently challenging due to the absence of a known ground truth. However, an indirect proxy for the true DelVG proportion can be derived from samples containing DelVG-specific mutations and/or full-length genome-specific mutations. These mutations produce mixed nucleotide signals at specific genomic positions, which can serve as a quantitative indicator of the relative abundance of the corresponding viral genomes within the sample.

We identified candidate mutations by analyzing the real datasets for variable positions (with several possible nucleotides), while systematically excluding regions between breakpoints, and the 150-nucleotide terminal segments of the genome. Using this approach, we identified 233 samples containing a total of 8,117 variable positions, with proportions likely attributable to either DelVGs or full-length genomes. Figure 8 presents the comparison of the DelVG proportions estimated by DIPScan to the observed mixed-position frequencies serving as a proxy for true proportions. The resulting Pearson correlation coefficient (*r* = 0.96, *p <* 2.2 *×* 10^−16^) indicates a high agreement between DIPScan’s estimates and the empirical proxy, supporting DIPScan’s accuracy in quantifying DelVG abundance.

**Fig 8.**
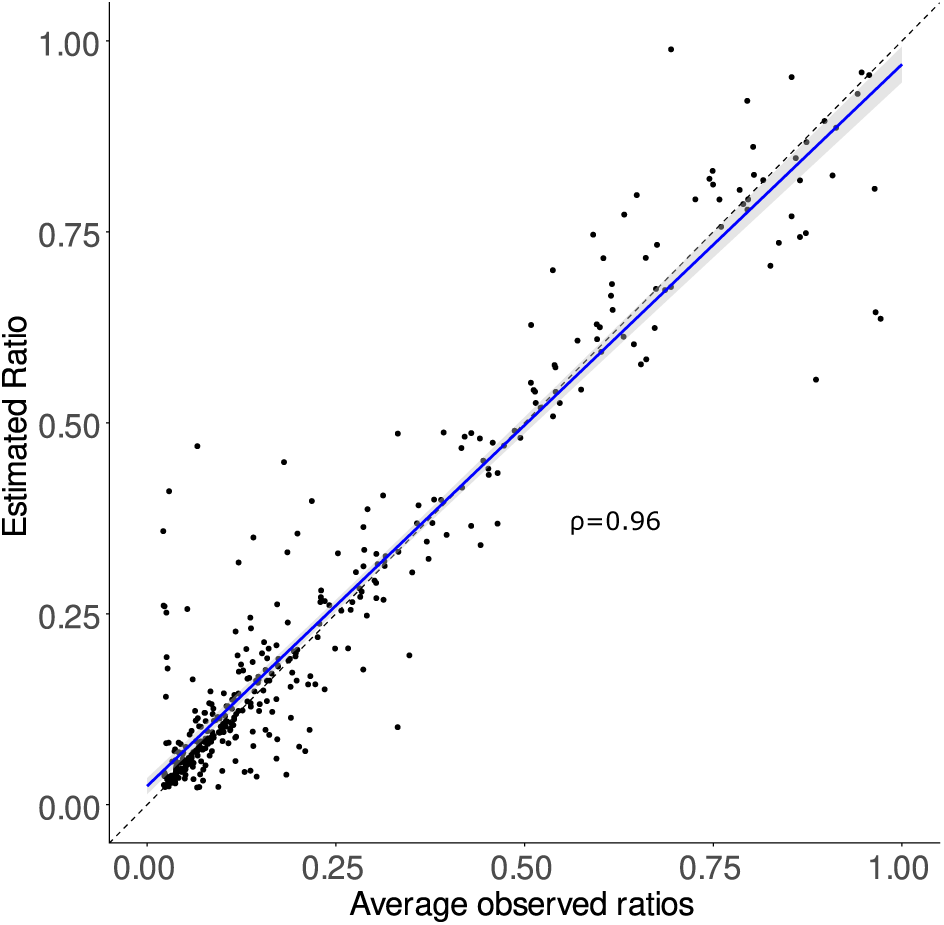
Evaluation of DIPScan proportion estimation on real datasets. Ratios between DelVG-specific coverage and full-length sequence coverage estimated by DIPScan (y-axis) are compared to the ratios derived from DelVG-specific (or full-length-specific) mutation frequencies at highly unambiguous positions (x-axis). Each point corresponds to the average over potentially multiple mutations for a given sample. The high correlation (0.96) between DIPScan estimates and ratios derived from unambiguous positions validates its use for downstream consensus correction.

#### Consensus correction

Across the whole dataset, 333/627 defective segments (53.1%) were subjected to correction of at least one position (41.4% for PB1, 25.5% for PB2, 32.1% for PA, 0.3% for HA, 0.3% for NA, and 0.3% for NP). Among the corrected positions, DIPScan changed the nucleotide in 17.7% of the cases. In the other cases (82.2%), DIPScan incorporated an ambiguous nucleotide.

To assess DIPScan’s accuracy in correcting the consensus sequences, a ground truth dataset was build consisting of 25 positions across 19 samples where nucleotides could be unambiguously (manually) attributed to either the full-length genome or the defective genome.

As depicted in Fig. 9, 24/25 positions (96%) in the selected ground truth dataset were correclty handled: 4 positions correctly changed, 13 positions correctly retained (13 unchanged), 7 positions appropriately masked, and only one position incorrectly changed, corresponding to a difficult case of DelVG proportion estimation.

**Fig 9.**
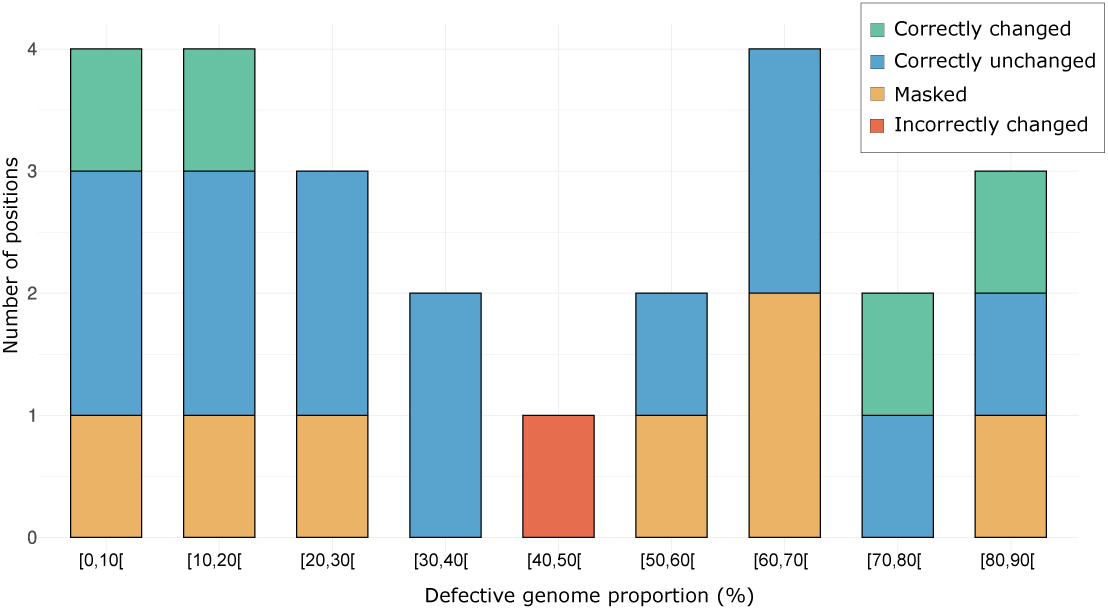
Validation of consensus correction on real datasets. Validation of consensus correction using 25 unambiguous mutations from 19 real Influenza virus datasets. Each of the 25 positions is classified by : (i) The proportion (estimated by DIPScan) of the detected DelVG covering the position (x-axis) and (ii) the outcome of the consensus correction: correctly changed (green), correctly unchanged (blue), masked with an ’N’ (orange), or incorrectly changed (red). The y-axis shows the number of position in each category. We observe that globally, most of the positions (96%) are correctly changed, unchanged or masked, and 1 position is incorrectly changed.

Masked positions resulted from: (i) Poor alignment (5 positions from 5 samples; detected via comparison of expected depth and coverage of the mutated position), (ii) Inability to match estimated full-length proportions to nucleotide frequencies (1 position from 1 sample), (iii) Estimated full-length proportions below the 2% noise threshold (1 position from 1 sample).

For samples with *>* 50% DelVG content, the unexpected cases of correctly unchanged positions occurred when the initial alignment (used to generate the consensus) was imperfect and resulted in the coincidental introduction of full-length-specific nucleotide into the consensus instead of the dominant DelVG-specific nucleotide.

#### Breakpoint location hotspots

With DIPScan results in hand, we could further characterize the DelVG sequences. In particular, we used the results from the 551 real dataset samples to examine the breakpoint positions along the genome and identify regions enriched with breakpoints (“hotspots”).

To do so, we grouped the results by virus subtype and segment, as they may exhibit distinct patterns of genomic deletions (see Fig. 10, S4 Fig).

**Fig 10.**
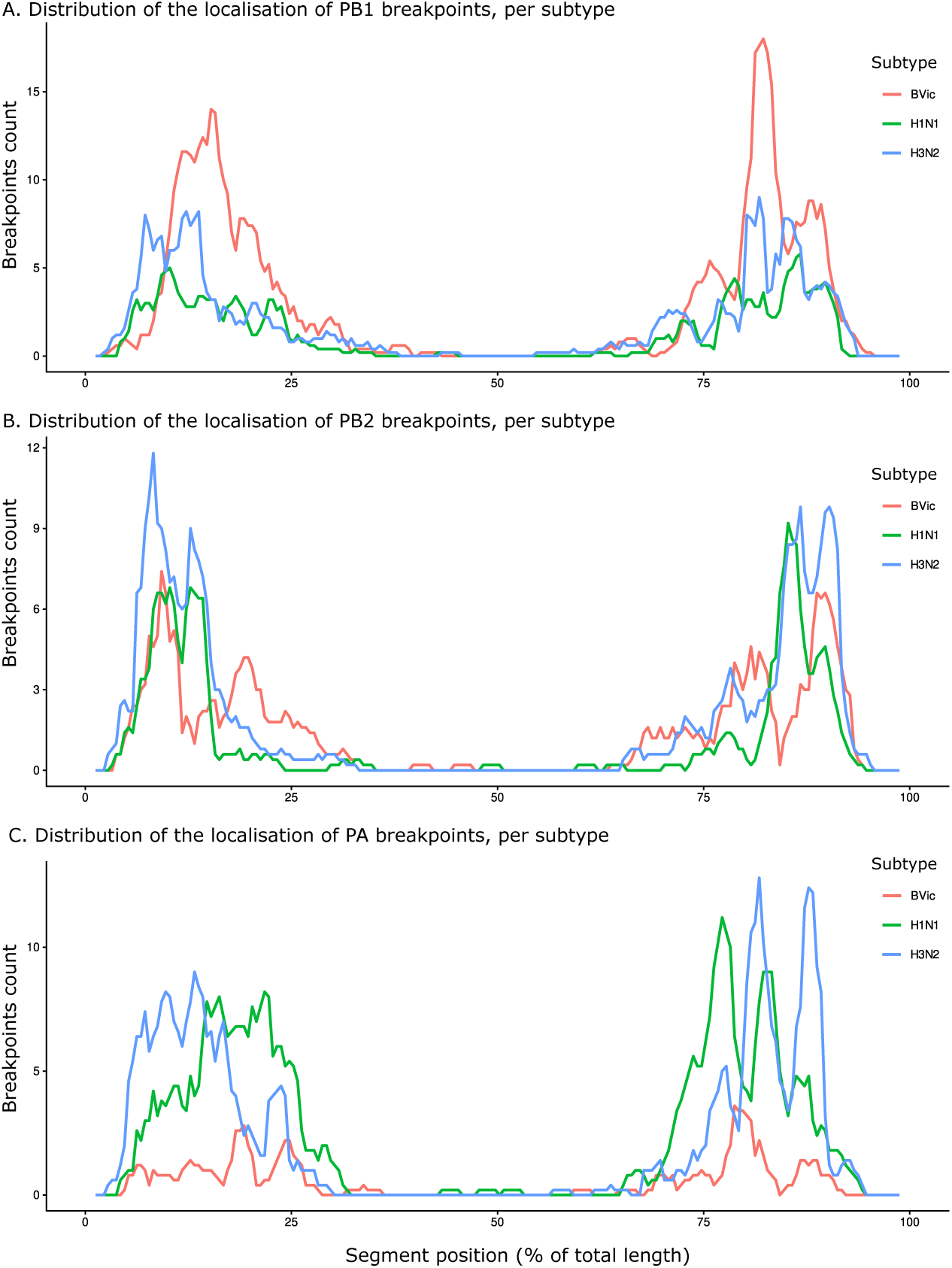
Analysis of potential breakpoint hotspots. The distribution of detected breakpoint ends (y-axis) is shown across the genomic segment, with the x-axis representing the location as a percentage of the segment’s total length. This distribution is colored by Influenza virus subtype of the 551 analyzed samples: B/Victoria in red, A/H1N1pdm in green, and A/H3N2 in blue. Only the 3 main affected segments are represented: **A.** PB1, **B.** PB2 and **C.** PA. This confirms that clear breakpoint hotspots exist near the segment ends of the three analyzed subtypes, and the three mainly affected segments (PB1, PB2, and PA).

Our analysis demonstrates that DelVG breakpoints are clearly not uniformly distributed across the viral genome (Fig. 10), instead exhibiting distinct ”hotspots” that vary by segment and subtype. Moreover, the breakpoint start-end locations seem to be dependent from each other (See S4 Fig). These observations align with previously reported patterns of DelVG breakpoint localization in influenza [45–47].

More specifically, breakpoint distribution patterns revealed subtype-specific variations in DelVG generation depending on the segments.

For the PB2 segment (Fig. 10.B), all three subtypes displayed 5’ breakpoints predominantly at around 9% of the genome length, and a second peak was observed around 12% for A/H1N1pdm and A/H3N2, while B/Victoria exhibited a shift to 19%. The 3’ breakpoints clustered around 89% for all three subtypes, and a second peak was observed around 85% for A/H1N1pdm and A/H3N2, while B/Victoria exhibited a shift to 80%.

For the PB1 segment (Fig. 10.A), A/H1N1pdm showed 5’ breakpoints at 10%, A/H3N2 at 8% and 13%, and B/Victoria at 15%. A/H1N1pdm and A/H3N2 showed 3’ breakpoints at 78% and 86%, B/Victoria also shared a peak at 86% and exhibited a second one with a shift to 88%.

In the PA segment (Fig. 10.C), A/H1N1pdm showed 5’ breakpoints between 16-22%, A/H3N2 between 7-16%, and B/Victoria exhibited two main peaks at 19% and 24%. All three subtypes displayed 3’ breakpoints at 87%, then A/H1N1pdm and A/H3N2 had other peaks at 77% and 82%, and B/Victoria at a second peak at 79%, highlighting subtype-specific variations in DelVG generation.

While confirming these breakpoint hotspots, these results further validate the ability of DIPScan to accurately detect DelVG sequences, and its usage in a routine setting.

## Discussion and perspectives

The widespread use of genome sequencing has enabled more precise and large-scale viral epidemic surveillance, for a broad range of pathogens, from EBOV [48], ZIKV [49], to SARS-CoV-2 [50]. While many virus epidemics are now monitored using molecular data, respiratory viruses remain a major concern, and generate the most massive amounts of data [51]. The scale of these datasets, combined with the need for automated consensus generation and curation to ensure accuracy, presents a significant bioinformatics challenge. Consequently, bioinformatics workflows have incorporated specialized functionalities for viruses, including reference selection, segmented genome support, and lineage assignment based on current nomenclatures.

Despite these advances, routine bioinformatics workflows have lacked the capability to account for the specific features of defective genomes when generating the consensus. To address this gap, we developed DIPScan, a Nextflow workflow designed to detect Deletion-containing Viral Genomes (DelVGs) that result from large deletions, and correct the consensus sequence when possible.

Using simulated data, we demonstrated that DIPScan accurately detects DelVGs (accuracy of 98.9%) and accurately estimates DelVGs/full length genome proportion (correlation of 0.99 between expected and estimated DelVG proportions). Additionally, DIPScan’s correction of consensus sequences is reliable, with 97.2% of the

DelVG-specific mutations being either replaced with the correct nucleotide present in full-length genomes or masked, into the final consensus sequence.

Analysis of 551 patient derived sequencing samples revealed a good agreement between DIPScan and manual curation, especially for predominant DelVGs (*>* 50%). Discrepancies were mostly due to manual errors. However, when DelVGs were detected only by DIPScan, they corresponded to genuine internal deletions or to complex cases requiring further review. In all scenarios, DIPScan enabled the rapid identification and correction of DelVGs at a large scale.

Furthermore, the automatic execution of DIPScan on real datasets identified clear preferential genomic breakpoints, “hotspots” positions for defective genomes, which is in accordance with previous reports [10, 19], and confirms it on a large number of samples.

In conclusion, DIPScan provides an accurate, efficient and scalable workflow for detecting DelVGs in influenza virus Illumina sequencing datasets, and, importantly, for correcting consensus sequences by accounting for DelVG-specific mutations.

The tool has been integrated into the NRC’s routine sequencing pipeline to screen for potential DelVGs. DIPScan is continuously being developed, and future developments will focus on two main areas.

**Extension to Other Pathogens**: We plan to test and benchmark DIPScan on datasets from other viruses known to produce DelVGs, such as RSV and SARS-CoV-2 [16]. This may require parameter optimization to ensure robust performance across different viral genomes.

**Detection of diverse types of defective genomes**: A key priority is to extend support to other defective genomes, including copy-back and rearrangements. While these are less frequent (although may explain the 29 influenza B samples identified as ”defective only in manual”) and present a greater challenge for defining genomic regions for correction, we could refine our methodology to identify them. One promising approach is to leverage the chimeric read detection capabilities of mappers like STAR, which could provide the necessary signal to detect these complex structural variants.

## Supporting information

**S1 Text. DIPScan workflow description.**

**S2 Text. Simulated dataset generation and analysis methods.**

**S3 Text. Real dataset analysis methods.**

**S1 Table. Description of simulated datasets.**

**S1 Fig. Illustrative example of the estimation of DIP + Full length sequences abundance.**

**S2 Fig. Estimated DIP Proportion (simulated data).**

**S3 Fig. Correction of common mutations (Simulated data). S4 Fig. Analysis of potential breakpoint pairs hotspots.**

## Implementation and data availability

DIPScan workflow is implemented in Nextflow [52], which makes it easily executable on many computing platforms, reproducible, and scalable. Each step uses software containers, which enables a high control of the software environment executed. DIPScan is freely available at https://github.com/pasteur-cnrvir/dipscan along with its documentation, and in Zenodo (DOI: 10.5281/zenodo.18816196).

The code and the datasets used to test DIPScan are available at https://github.com/pasteur-cnrvir/dipscan_analysis/ and Zenodo (DOI: 10.5281/zenodo.18816200).

Real patient derived datasets analyzed in this study are available in the NCBI Sequence Read Archive (SRA) under BioProject accession number PRJNA1426172. Simulated datasets are avaible on Zenodo (DOI:10.5281/zenodo.18802934).

## Author contributions statement

F.L. and M.R.W. conceived the project, F.L., K.D.S., N.N. and M.R.W. conducted the research and wrote and reviewed the manuscript.

## Acknowledgments

This work has been partially funded by the European Union’s Seq4Epi project (Grant Agreement Project 101113174 - SEQ4EPI EU4H-2022-DGA-MS-IBA-1).

